# Selectively expressed RNA molecules: a new dimension in functionalized cell targeting

**DOI:** 10.1101/2022.05.10.491309

**Authors:** Marco Hoffmann, Sylvie Schüle, Christina Hoffmann, Frederik Rastfeld, Sven Gerlach, N. Hersch, Helene L. Walter, Dirk Wiedermann, Gereon R. Fink, Rudolf Merkel, Heribert Bohlen, Maria A. Rueger, Bernd Hoffmann

**Author notes:** Address correspondence to: Dr. Bernd Hoffmann: Institute of Biological Information Processing, IBI-2: Mechanobiology, Forschungszentrum Jülich, 52425 Jülich, Germany.

## Abstract

Exact targeting of specific mammalian cell types or diseased cells is one of the most urgently needed prerequisites for a new generation of potent pharmaceuticals. Different approaches have been pursued, failing mainly due to a lack of specific surface markers in most cases. Developing a completely novel RNA-based methodology, we can now ensure exact cell targeting and simultaneously combine this with selective expression of effector proteins, thereby functionalization of the target cell for therapy, diagnostics or cell steering. The specific combination of the molecular properties of antisense technology and mRNA therapy with functional RNA secondary structures allowed us to develop selectively expressed RNA molecules for medical applications. These so-called seRNAs remain inactive in non-target cells and are only activated by partial degradation to induce translation in preselected cell types of interest. Cell type specificity and type of functionalization are easily adaptable based on a simple modular system. In proof of concept in vitro and in vivo studies we used seRNAs as a highly selective platform technology for powerful glioblastoma cancer cell targeting and significantly reduce brain tumors of mice without detectable side effects with just a single treatment within days. Our data open up new potential avenues for the efficient treatment of various cancers and other human diseases.

## Introduction

Fundamentally all modern medical treatments against cancer, as for viral, metabolic and genetic diseases, would massively benefit if the affected cell could be specifically identified and targeted. Such ideal medication would be locally effective in the optimal functional concentrations without causing significant side effects in the surrounding tissue or the entire organism.

For decades, approaches for targeted delivery of therapeutics in cancer cells have existed and typically involve systemic administration of therapeutics packaged in nanocarriers (NCs) or localized delivery to the diseased tissue. Encapsulation of therapeutic molecules (e.g., small molecule inhibitors, chemotherapeutics, siRNA) in NCs can improve their solubility and bioavailability, alters their bio-distribution, and facilitate their entry into the target cell. “Passively” targeted NCs, which utilize the enhanced permeability and retention effect^1^ of many solid tumors and subsequently depend on endosomal uptake, are the most extensively explored strategy for targeting cancer systemically^2^. Multiple passively targeted NCs have been approved over the past 20 years for medical applications^3^. However, due to significant uptake also by healthy cells, side effects remain high, limiting their medical use. This essential drawback becomes even more important since a significant proportion of such passively targeted NCs is often lost upon in vivo application due to non-specific surface binding or clearance by the mononuclear phagocytic system^4^.

Active cellular targeting was developed as a complementary strategy for improving tumor localization of NCs by increasing their targeting efficiency and enhancing retention and uptake at the target site^5, 6^. However, although having ultimate potential, active targeting largely depends on overexpressed or cell type specific surface markers of diseased tissues or cells and their ability to induce internalization upon specific NC surface binding^4, 7^. One of the currently most promising targeting approaches is the use of Car-T-cells as effective medication against B-cell leukemia and others^8–10^, although the underlying targeted surface marker is not a disease-associated protein but the B-cell specific CD19 surface protein. Therefore, the therapeutic application is always accompanied by a severe B-cell aplasia, which massively affects healthy B-cells upon treatment, limiting the general applicability of the method^11^. Consequently, lack of diseased cell-specific markers combined with further challenges at the level of premature release of therapeutics^2^, low endosomal release rates^12^ and effective direct exocytosis of internalized NCs^13^ resulted in only a small number of clinical trials that are based on actively targeted NCs^14^. Furthermore, targeting approaches of just a single cell-surface receptor on tumor cells disregards tumor heterogeneity and promotes selection toward the survival of resistant clones^15, 16^.

Interestingly, proteome and transcriptome analyses suppose that suitable surface markers will remain the major challenging limitation for classical active targeting approaches. The same analyses, however, revealed a much higher probability of identifying characteristic molecules on RNA level in diseased cells, irrespective of their underlying function (e.g. mRNA, (l)ncRNA)^17–19^ and subsequent protein localization. This problem is exemplified by glioblastoma brain tumors, in which several surface markers show increased expression but never true specificity^20^, whereas a large number of glioblastoma-specific RNA molecules has been identified at the RNA level^21^. Transferring cell targeting to the RNA level within the cytoplasm would therefore open up a formerly unprecedented range of new targets and application possibilities.

Here, we have developed a completely new system of selectively expressed RNA molecules (seRNA) for efficient targeting and simultaneous functionalization of any cell type. Based on nucleic acids that can be transferred as DNA or RNA in a non-targeted manner, resulting RNA molecules encode for an effector protein but remain inactive in healthy cells while being activated and translated in predefined target cells only. Targeting of diseased cells is therefore transferred from the outside of the cell to the immense intracellular pool of disease specific molecules. This targeting technology is made possible by a unique combination of various highly conserved RNA-based regulatory mechanisms^22–30^. The optimal arrangement of all individual domains to each other in a simple modular system guarantees for translational inactivity in non-target cells. In contrast, target cells induce partial degradation of the same seRNA molecules by sense-antisense interactions, whereby Internal Ribosomal Entry site (IRES)-blocking sequences are removed to enable IRES-dependent translation while simultaneously inhibiting further degradation. Consequently, the developed seRNAs represent a completely new and powerful platform technology for cell targeting and functionalization to open new routes in biotechnological and medical applications. Rigorous testing of specifically developed seRNA molecules as drug candidates against glioblastoma proof cancer cell specific activation and allow for efficient treatment of brain tumors in mice within days.

## Results

### A multidomain structure enables unique cell targeting properties of seRNA molecules

seRNA molecules comprise all classical domains known from mRNAs (Fig. 1A, domains 1 + 2 as well as 7 to 9). Their target specific translational activity, however, depends on various additional sequence motifs. Most importantly, an antisense sequence that is complementary to freely selectable target RNA (Fig. 1A, domain 3) is positioned as part of an untranslated 5’-region. This sequence additionally must contain small upstream open reading frames (uORFs). These uORFs partially remove CAP-dependent ribosomal subunits from the seRNA molecule and inhibit expression of the seRNA encoded effector protein by overlapping with the main reading frame (domain 7). Such regulatory mechanisms have been known for decades and are present in various mRNAs with starvation induced translational activation^26, 30^. Since CAP-dependent translation is omitted for seRNAs, an Internal Ribosomal Entry Site (IRES) (Fig. 1, domain 6) is responsible for translational initiation of the effector ORF. However, such a construct would translate effector ORFs (Fig. 1, domain 7) constitutively and independent of cell type. Fortunately, IRES functionality is primarily based on secondary structure formation and already minor alterations in structure entirely block translational initiation activity^29^. We utilize this structural sensitivity here by incorporating short RNA-sequences 5′ to the IRES that interfere with complementary IRES sequences to induce secondary structure rearrangements that function as IRES-blocker (Fig. 1A, domain 4). Having such an incomplete seRNA construct in hand, IRES would remain inactive in non-target cell types but sense-antisense interaction would induce unwanted degradation of the complete seRNA molecule in target cells. We therefore use viral RNA sequence motifs that efficiently block 3′-directed RNA exonuclease activity^25^. This results in protection from further degradation of downstream located seRNA sequences. By placing such a motif between IRES-blocker and IRES (Fig. 1, domain 5), seRNA molecules display their full potential with blocked IRES activity in non-target, i.e. healthy cells. In target cells, however, the formation of double stranded RNA motifs induces seRNA degradation, removing the IRES-blocker but leaving downstream IRES and effector coding sequences unaffected to induce targeted translation and cell functionalization.

**Figure 1:**
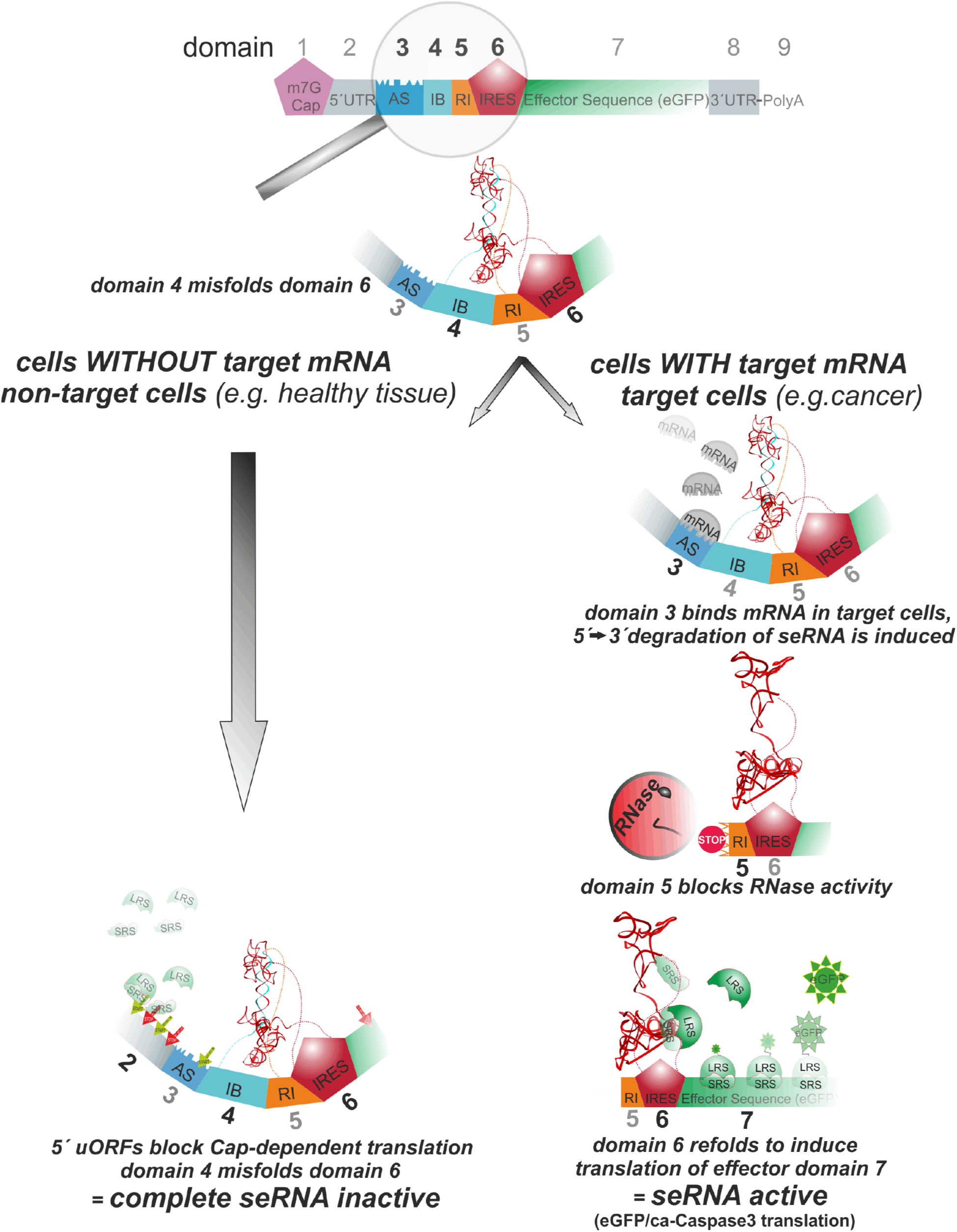
seRNA functional mechanism. Based on specific sense-antisense interactions, seRNA molecules induce translational activity. Upon transfer of seRNAs into non-target cells missing target mRNAs (e.g. cancer marker) (left), seRNAs remain non-functional due to structural misfolding and CAP induced ribosomal silencing by uORFs (green and red arrows). In contrast, target cells (e.g. cancer cells) induce partial seRNA degradation due to sense-antisense interaction (right). Intrinsic block of degradation induces functional refolding of seRNA to allow IRES-dependent efficient translation of effector sequences in target cells only. UTR = untranslated region, AS = antisense domain, IB = IRES blocker, RI = RNase inhibitor

### seRNA molecules specifically target and functionalize pre-chosen cell types

To verify functionality, seRNA constructs were generated to specifically target glioblastoma cancer (U87) cells using a conserved keratin motif as antisense sequence and eGFP as effector. To prove specificity on a broader range of cell lines, breast cancer (MCF-7) cells were also chosen. The expression of keratin as an established diagnostic cancer marker was confirmed by immunocytochemistry and qRT-PCR. Primary human foreskin fibroblasts and primary cortical neurons served as healthy non-target control cells without detectable keratin expression (Fig. 2a+b).

**Figure 2:**
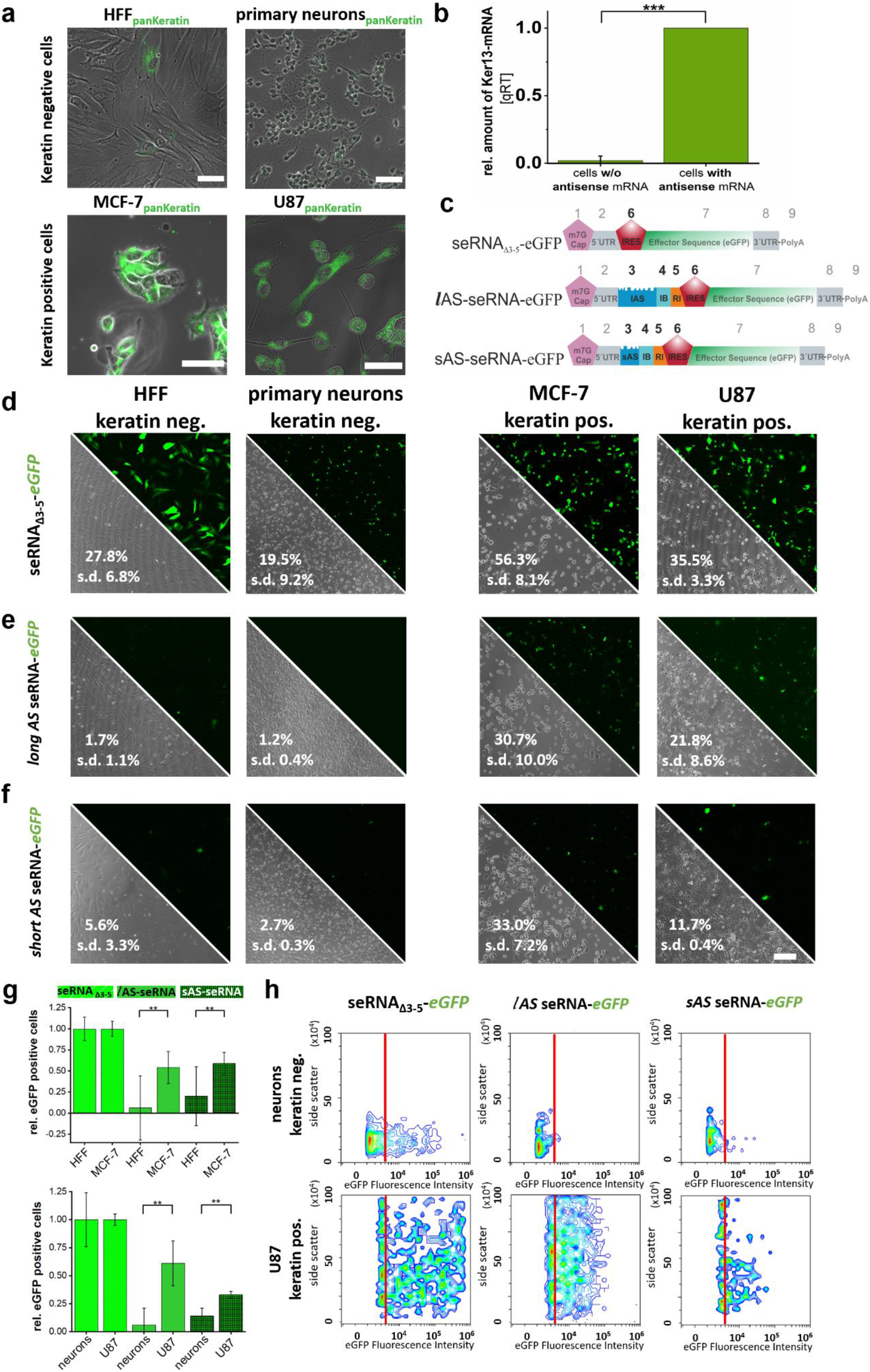
seRNA targeting specificity. Using keratin as selectively expressed target for seRNA activation, keratin negative HFF and primary neurons as well as keratin positive cancer cell lines MCF-7 and U87 were stained (a). Normalized keratin signal intensities are indicated in (b). Domain architecture of non-selective and selectively expressed RNA constructs is indicated (c). Constructs were used for expression analysis in non-target and target cells after transfer as DNA-expression plasmid (d to f). eGFP positive cells were either given as percentage of all cells (d to f) or as the relative value using transfection efficiencies of non-selectively expressed RNA constructs as reference (g). Flow-cytometry analyses of primary cortical neurons (top) and U87 glioblastoma cells expressing non-selective and seRNA constructs. Threshold (red line) indicates 99% of all cells in an untransfected control (h). Scale bar = 50 μm. n = at least 3 independent experiments.

Transfer of seRNA expression plasmids lacking regulatory targeting domains 3 to 5 (seRNA_Δ3-5_-eGFP) (Fig. 2c) proved eGFP expression in all cell types with transfection efficiencies ranging from approximately 20% for cortical neurons to 56% for MCF-7 cancer cells (Fig. 2d).

In contrast, upon transfer of all-domain containing seRNAs (Fig. 2c), eGFP expression was blocked almost completely in keratin-free HFF and primary cortical neurons (Fig. 2e). However, the same constructs induced stable eGFP expression in about 60% of all transfected cells in the presence of keratin in MCF-7 and U87 cells (Fig. 2e+g). The inhibitory effect of a changed number of 5′ uORFs was analyzed by shortening the antisense sequence from 615 bp to 150 bp. This reduced the number of uORFs from 5 to 2 while keeping the last uORF overlapping with the main ORF intact. Cancer cell targeting and specific translational activation of seRNAs remained functional (Fig. 2f+g). Also overall translational repression for both seRNA constructs was active in non-target cells. However, low base level activity in non-target cells was approximately doubled upon uORF reduction. Flow cytometry analyses confirmed these results with inactive seRNAs in non-target primary neurons but effective cancer cell targeting and induced expression of the effector (eGFP) in U87 glioblastoma cells. As shown before, seRNAs with long antisense domain were slightly more effective than short ones. Furthermore, high CAP-dependent overexpression of eGFP in seRNA_Δ3-5_-eGFP constructs was missing for all full-length seRNAs with more homogeneous expression levels in target cells (Fig. 2h).

### seRNAs are activated by partial degradation of dsRNA sequences in target cells but do not induce cytokine response

Functional analyses of seRNA translational activation in target cells were performed for long antisense seRNA expression constructs with GFP as effector after transfer into target (MCF-7) and non-target cells (HFF). Using seRNA-specific probes for sequences upstream and downstream of domain 5 (RNase inhibiting sequence) in simultaneous two-probe qRT-PCR analyses, stable levels of full length seRNA molecules in non-target cells were detected over time. Here, seRNA concentrations increased with time to reach steady state levels after approximately 8 hours (Fig. 3a and b), proving expression of full length seRNAs in non-target cells and translational inactivity due to seRNA self-inhibition. Concentrations of seRNA 5’-regions and 3’-regions show an almost identical concentration behavior, which argues for molecular stability regulation as a whole, as one would expect for mRNA molecules. In contrast, upon seRNA transfer into target cells, upstream and downstream probes identified significantly diverging concentrations of underlying templates. While concentrations of seRNA 3’-region increased with time to similar steady state levels after approximately 8 hours as shown for non-target cells, concentrations of seRNA 5’ region barely increased with time to steady-state levels of clearly less than 1% of the corresponding 3’ region. These data clearly prove massively reduced lifetimes for 5’-regions upstream of domain 5 and simultaneous stabilization of remaining 3’-seRNA fragments in target cells. Blocking RNaseH1 activity we confirmed its partial influence in recognition and digestion of formed dsRNA domains upon expression of seRNAs in target cells. Here, presence of inhibitor enhanced concentration of undigested, full-length seRNA and therefore increased amounts of 5’regions in two-probe qRT-PCR analyses (Fig. 3c).

**Figure 3:**
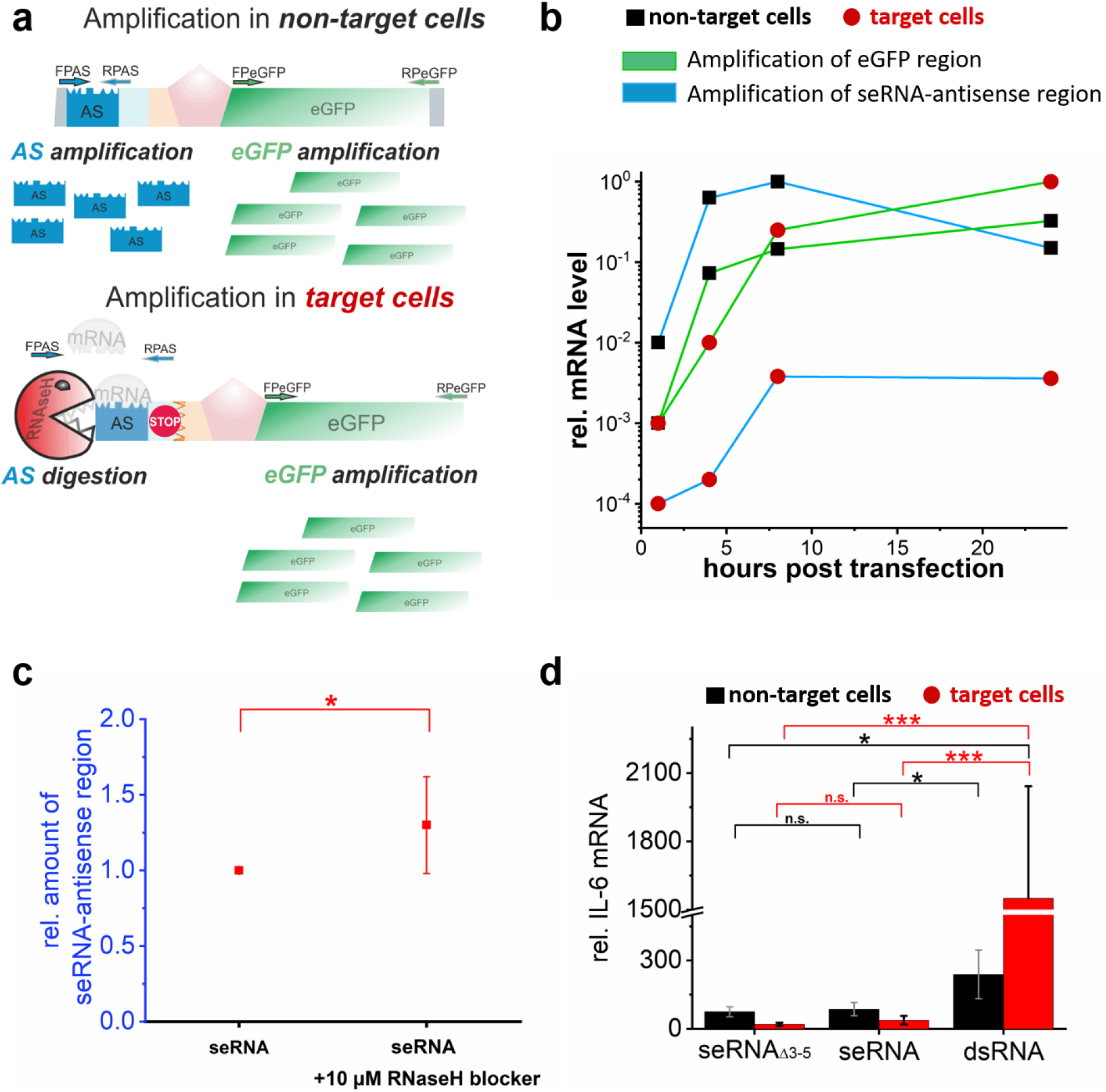
seRNA functional characterization. Using two different probes in parallel (a), qRT-PCR experiments were performed after the transfer of seRNA-eGFP in non-target and target cells for indicated time periods (b). Inhibition of RNase HI in non-target and target cells after transfer of seRNA. 8 h after transfection two-probe qRT-PCR analyses were performed (c). Cytokine response analysis after transfer of stable double-stranded RNA (dsRNA), full-length (seRNA) and shortened seRNA_Δ3-5_-eGFP constructs into non-target and target cells. 24 h after transfer, IL-6 expression levels were quantified by qRT-PCR (d). All values are given with s.d. n=3 or more.

Double stranded RNA molecules have been shown to induce toll-like and RIG-I-like receptor dependent innate cytokine responses in vitro and in vivo^31, 32^ with potentially massive local or systemic inflammatory responses, which conflict with medical applications. Since seRNA activation also depends on temporal intracellular RNA double-strand formation and recognition, we tested for cytokine expression after transfer of seRNA expressing plasmids into target and non-target cells. Stable double-stranded RNA molecules of comparable length as used as antisense in seRNA molecules were taken as positive controls. Such stable dsRNAs induced massive IL-6 expression in non-target and target cells by a factor of up to 1350 compared to seRNA_Δ3-5_ within 24 h (Fig. 3e). In contrast, for full-length seRNAs no antisense-induced cytokine induction could be detected. Regardless of whether constructs were expressed in non-target or target cells, antisense-specific IL-6 expression remained low with even lower values in target cells than non-target cultures.

### IRES-blocker positioning facilitates an additional level of translational regulation

We propose that quiescence of seRNA molecules in non-target cells as well as putative leakiness levels primarily depend on the ability to effectively interfere with the IRES secondary structure in full-length seRNA constructs. At the same time, this block in IRES activity must not hamper IRES functional refolding upon seRNA activation by partial degradation in target cells. We therefore analyzed IRES-blocker (IB) specificity (domain 4) in more detail and generated constructs with IRES complementary sites to interfere with largely unfolded IRES regions (IB1), stem-loops (IB2), and stem segments (IB3) of the IRES secondary structure. In addition, IB3 was located at the central position (pseudo knot) while IB2 affected a stem region with just supportive function^23^ (Fig. 4c to e). Secondary structure predictions for the IRES domain itself separated the pseudoknot region from surrounding IRES domains based on minimal free energy driven secondary structure predictions (Fig. 4a). Interestingly, the same algorithms indicate that addition of just the eGFP coding sequence changes structures of supportive IRES domains while the pseudoknot remains largely unaffected (Fig. 4b). However, and as expected, the secondary structure of the underlying seRNA_Δ3-5_ construct did not hamper untargeted eGFP translation in non-target (neurons) or target cells (U87). For full-length seRNA sequences with IRES blocker sequences IB1 to IB3 (Fig. 4c to e), stable interactions between IRES blocker and its binding site within the IRES were predicted in all cases. These interactions changed IRES secondary structures not only for the blocker binding domain but also for surrounding sequences. Most obvious structural adaptations were induced by IB3 with an elongated pseudoknot and largely aggregated supportive IRES domains.

**Figure 4:**
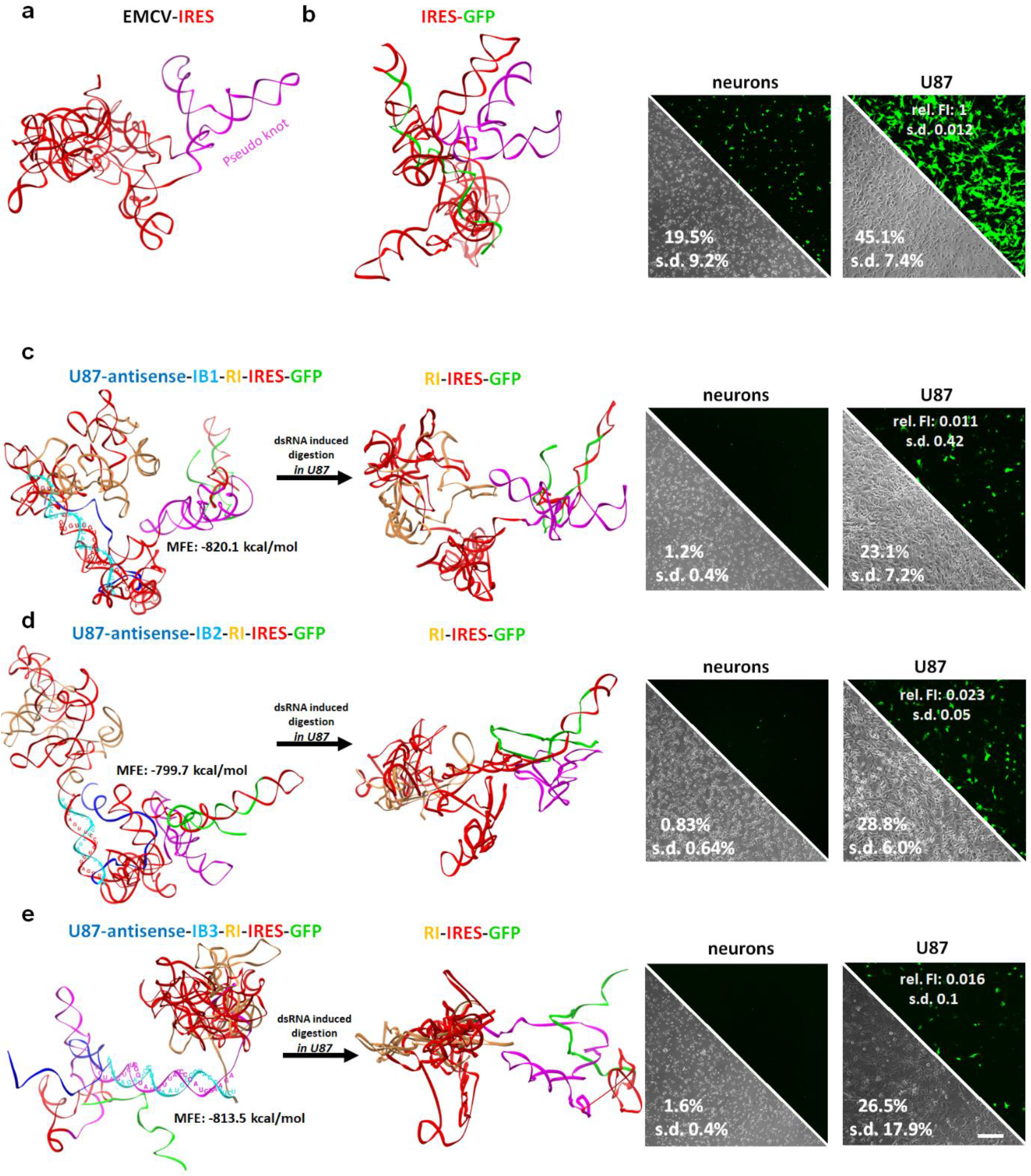
seRNA secondary structures. Based on minimal free energy (MFE) algorithms (RNAfold WebServer), secondary structure predictions were performed for the IRES only (a) and in the presence of eGFP coding sequence (b) as part of seRNA_Δ3-5_. Its GFP expression level is indicated for neurons and U87 glioblastoma cells (b). Secondary structures for seRNA domains 4 to 6 including few nucleotides of domains 3 (antisense) and 7 (GFP) with three different IRES-blocker sequences (IB1 to IB3, c to e) are given before (left) and after partial degradation in U87 glioblastoma target cells (middle). For all constructs MFE values are indicated. Their eGFP expression levels in non-target cells (neurons) and target cells (U87) are shown (green signal, right). Fluorescence intensities (relative units) and quantitative flow cytometry data (in %) are indicated for IB1-3 in each image. Unaffected cell morphology is indicated in phase contrast. All values are given with s.d. Scale bar = 200 μm. n=3 or more.

All full-length seRNA constructs were blocked in functional analyses to induce eGFP expression in non-target primary neurons (Fig. 4c to e). In U87 target cells, however, all constructs facilitated eGFP expression predictively due to selective RNA degradation with subsequent loss of IB1 to IB3. IRES secondary structures differed despite IRES-dependent eGFP expression intensities being comparable for all shortened seRNA constructs. Assuming comparable transfection efficiencies for all full-length seRNA and the constitutively active seRNA_Δ3-5_-eGFP construct, data show for approximately 50-65% of all seRNA transfected cells efficient IRES refolding with subsequent induction of eGFP expression. Interestingly, the data indicate that pseudoknot secondary structure perturbations of IB3 impair functional IRES refolding more than IB1 and IB2 through high standard deviations in the achieved eGFP-efficiencies. These data argue that the best target-specific activation is possible using blocker sequences that induce just minor and spatially restricted IRES secondary structure misfolding.

### seRNA-based glioblastoma cell targeting combines optimal specificity with effectively induced cancer cell death

Due to extraordinary seRNA targeting abilities with functional activation only in target cells, we replaced eGFP with a constitutively active caspase 3 (ca-Caspase3). Caspase containing control constructs lacking the inner regulatory domains (seRNA_Δ3-5_–ca-Caspase3) proved this ability in every cell type tested. Here, approximately 90% of all transfected target and non-target cells, based on transfer of comparable eGFP-constructs, died in the presence of ca-Caspase3 within 24 h (Fig. 5a). In contrast, full-length seRNA-ca-Caspase3 constructs were entirely inactive in non-target primary neurons while equally effective as non-selective seRNA_Δ3-5_ constructs in U87 human glioblastoma target cells (Fig. 5b). Prolonged seRNA-eGFP expression for 48 h in combination with repeated seRNA delivery further enhanced transfection efficiencies to 95% in U87 cells and approximately 70% in primary neurons. Upon use of seRNA-ca-Caspase3, the same conditions resulted in total values of 60% apoptotic cells after 48 h while primary neurons remained thoroughly unaffected (Fig. 5c), therefore proving absent leakiness of seRNAs in non-target cells.

**Figure 5:**
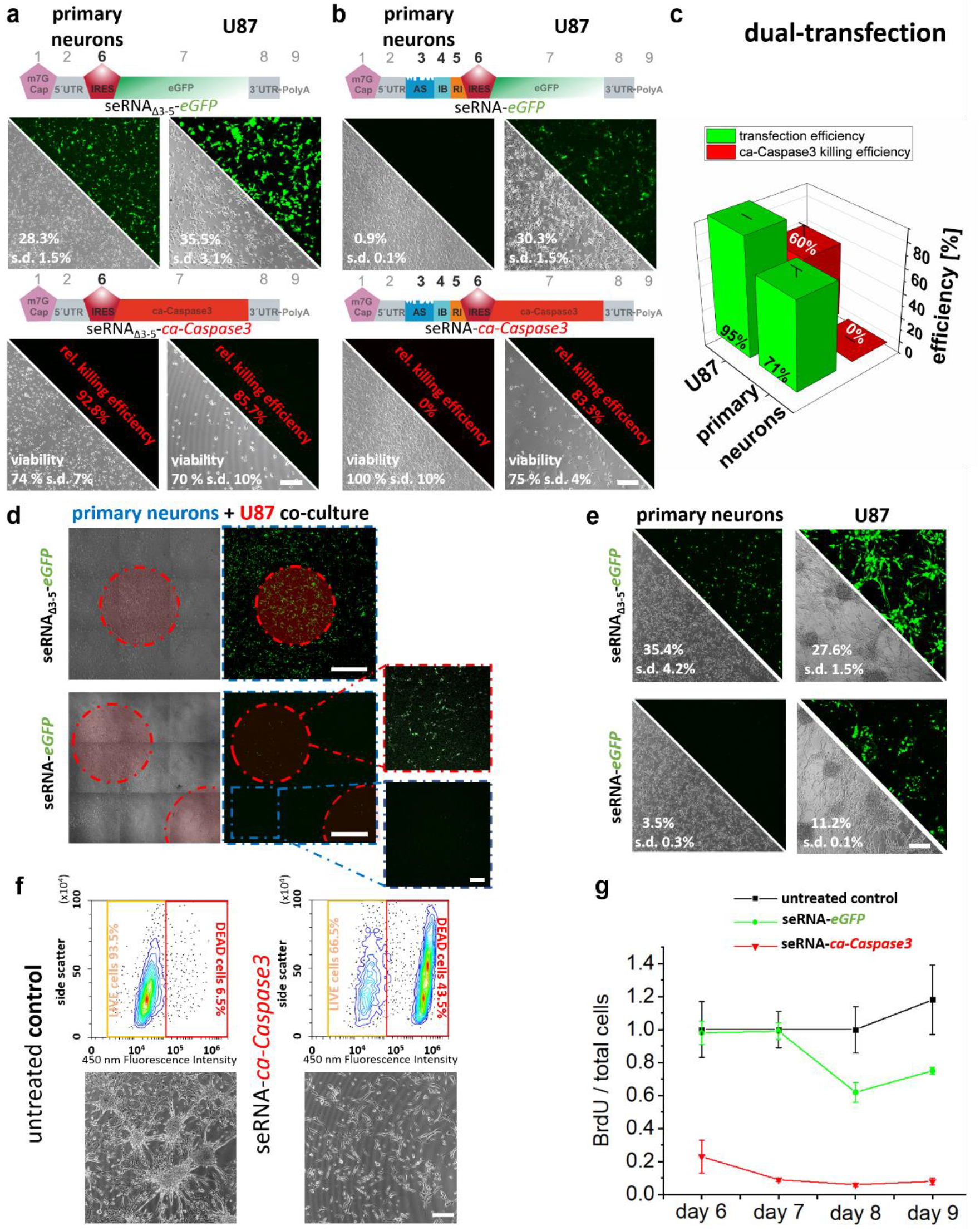
seRNA as a treatment approach against cancer. seRNA_Δ3-5_ (a) and seRNA (b) constructs with either eGFP or ca-Caspase3 were expressed in indicated cell types. Target specific expression of eGFP (green) and cell viability was analyzed by light microscopy and flow cytometry, given in %. Dual transfer at 24 h intervals of seRNA-eGFP and seRNA-ca-Caspase3 into the target and non-target cells with subsequent flow cytometry analysis 24 h after second transfer (c). Growth of patterned U87 cell clusters (red circles) in surrounded monolayer of primary neurons with subsequent transfer of seRNA_Δ3-5_-eGFP and seRNA-eGFP. For better visualization, zoom-in images are contrast-enhanced (d). Transfer of the same seRNA constructs into neuronal networks and U87 tumor-spheres after growth for 5 days (e). Single transfer of seRNA-eGFP and seRNA-ca-Caspase3 into U87 target cells 24 h after seeding during tumor-sphere formation. Spheres are visualized in comparison to untreated U87 cells in phase contrast. Caspase activity was characterized by flow cytometry. Enhanced fluorescence at 450 nm corresponds to reduced viability (f). Repetitive delivery of seRNA-eGFP and seRNA-ca-Caspase3 into young U87 tumor spheres at day 2, 4 and 6 in the presence of BrdU. Subsequent cell proliferation was analyzed at indicated time points (g). All values are given with s.d. Scale bar = 200 μm. n=3 or more.

To mimic glioblastoma in brain environment, neuronal networks from healthy primary neurons were formed after U87 cell growth for 3 days on the same plate. A single treatment with seRNA_Δ3-5_-eGFP proved effective transfer into both U87 tumors and neuronal networks with subsequent non-targeted GFP expression. In contrast, transfer of full-length seRNA-GFP confirmed also in mixed cultures the activation of effector expression only in U87 target cells and activity inhibition in non-target primary neurons (Fig. 5d).

Artificially formed solid U87 tumor spheres proved that the same target cell specific activation of full-length seRNAs remained as functional as found on single cell level (Fig. 5e). The same tumor spheres were used to analyze the effectiveness of seRNA activation and subsequent specific caspase expression. While such glioblastoma like spheres formed within 5 days in the absence of seRNA-ca-Caspase3, the formation was entirely suppressed after a single transfer of seRNA-ca-Caspase3 24 h after cell seeding (Fig. 5f). Data were also confirmed by quantitative flow cytometry analyses that showed massively induced cell death after transfer of seRNA-ca-Caspase3 (Fig. 5f). Furthermore, reduction of U87 cell proliferation in already formed tumor spheres by repetitive treatment with seRNA after 2, 4, and 6 days was analyzed by BrdU in long term experiments. While U87 cell proliferation remained stable on a high level in untreated cells, transfer of seRNA-eGFP controls proved continuous proliferation. In contrast, transfer of seRNA-ca-Caspase3 very effectively blocked proliferation by 75% to more than 90% throughout the whole analysis time (Fig. 5g).

### Therapeutic seRNA effectively treats glioblastoma in mice without detectable side effects

seRNA functionality was further verified in vivo in immunodeficient mice. Human U87 glioblastoma cells were injected into the striatum and grown for two days to allow for robust and mitotically active brain tumor formation (Fig. 6a, c). Treatments in independent control groups with either isotonic NaCl solution or full-length seRNA-GFP expression plasmids by stereotactic injection directly into the tumor showed highly comparable results without affecting tumor growth over time. In contrast, injection of the therapeutic construct seRNA-ca-Caspase3 into glioblastoma visibly reduced the tumor size as assessed by non-invasive MRI 14 days after treatment (Fig. 1b). Monitoring tumor development over time by MRI revealed continuous tumor shrinkage induced by caspase-expressing seRNA (Fig. 1e; p = 0,0001 and F = 17,58) as compared to tumors treated with control constructs (Fig. 1d p = 0,4078 and F = 0,9243). Likewise, sizes of caspase-seRNA treated tumors expressed in relation to control tumor volumes on day three, were significantly reduced (Fig. 1f; caspase: p = 0,0001 and F = 17,58; control: p = 0,4078 and F = 0,9243). Detailed daily monitoring showed no cytotoxic effects or behavioral changes in the treated animals at any time point, proving effectiveness of seRNA as therapeutic drugs in mice.

**Figure 6:**
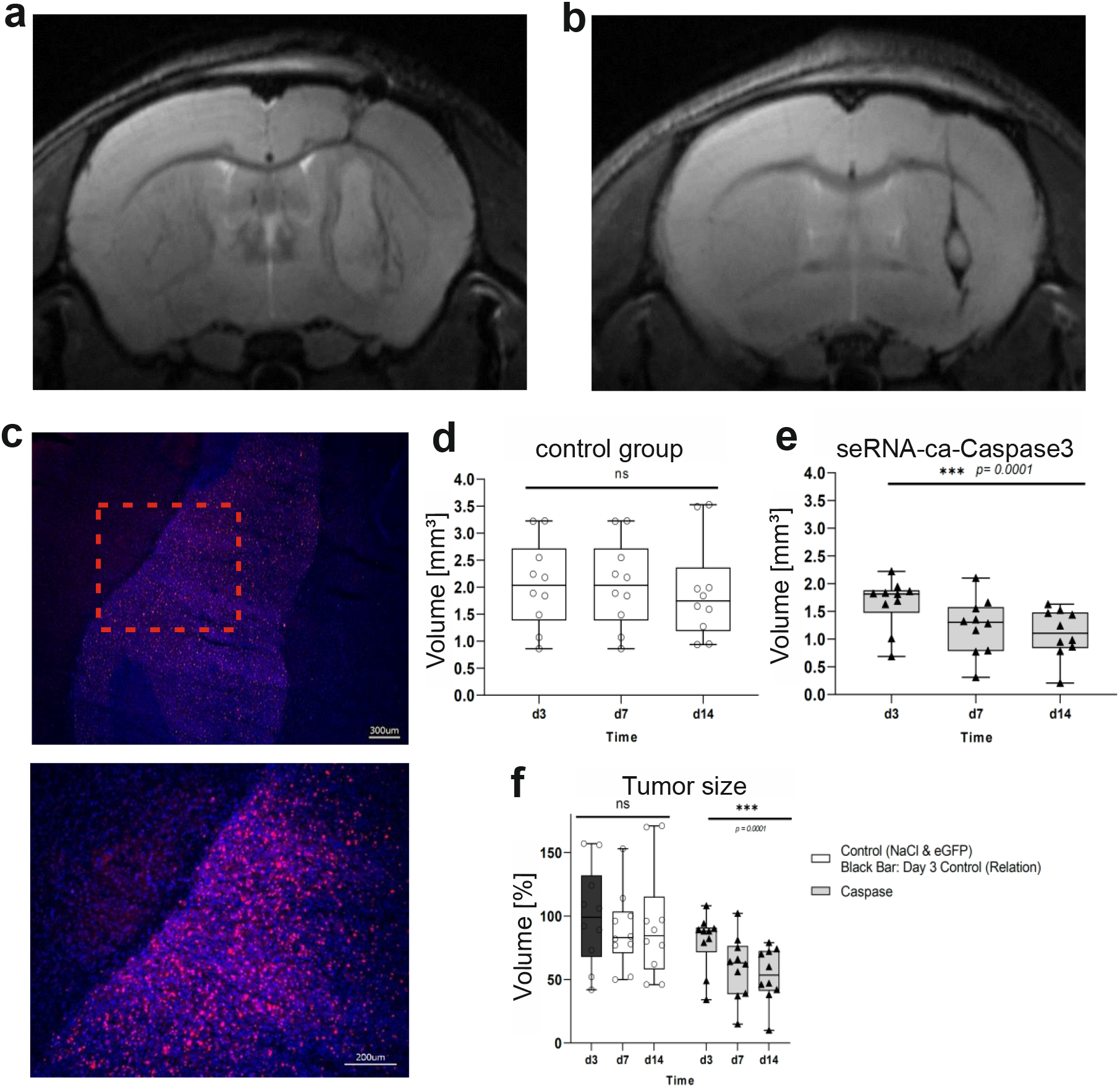
seRNA treatment of glioblastoma mice. Tumors of human U87 glioblastoma cells were grown in immunodeficient mice and analyzed by MRI 14 days after treatment with a therapeutically inactive seRNA-eGFP construct as control (a) or the therapeutic seRNA-ca-Caspase3 construct (b). Tumor formation and mitotic activity of tumor cells were verified by Ki67 immunoreactivity (red) and nuclear staining (blue) 16 days after tumor implantation (scale bar 300 μm) (c). A zoom in of the red dotted square is indicated below and reveals densely packed Ki67 positive tumor cells (scale bar 200 μm). Based on MRI images, tumor volumes were evaluated after treatment with control (seRNA-eGFP or NaCl) (d) or therapeutic seRNA-ca-Caspase3 constructs (e) at indicated time points. Statistical analyses were performed with a repeated measures ANOVA over the factor time with p = 0,5035; F= 0,4856 for (d) and p = 0.0001; F= 17,45 for (e). For direct comparison of control and therapeutic animals relative tumor volumes in relation to day 3 (d3) control mice are shown. Analysis via RM one-way ANOVA over the factor time (p = 0.0001) (f). For all statistical analyses box plots provide all single values for every time point.

## Conclusions

Techniques for efficient and freely adaptable cell targeting constitute one of the most urgent needs in biotechnology and medicine. Many different approaches have been developed, most of them depending on cell-type-specific surface markers^33, 34^, which are, unfortunately, most often missing. With seRNAs we have, for the very first time, developed a unique technology platform that allows cell targeting on RNA-expression level by shifting recognition from the outside to the inside of the cell, as known for siRNAs, and by combining this type of targeting with cell type specific translation of freely selectable effector proteins.

Current seRNA molecules are based entirely on recombination of known sequence motifs to combine their properties into a new mode of functionality. They bring together cell-type specificity of antisense RNA or siRNA approaches with cell functionalization abilities of mRNA techniques. This is made possible by regulated partial degradation in target cells only, which results in translational induction at IRES-dependent secondary sites. The Encephalomyocarditis virus (EMCV)-IRES used here for this purpose is well analyzed and functionally characterized by a homogeneous translation efficiency and independence from cofactors^23^. EMCV-IRES efficiency is typically lower than CAP-induced translation^35^ which confirms our expression data for seRNAs. We assume that EMCV-IRES can be easily replaced by other viral or even human IRES sequence motifs to thus add further IRES-specific functionalities to seRNA molecules, such as different translation efficiencies or cell type-specific regulation by binding proteins^36^.

The efficient blocking of IRES activity in full-length seRNA is crucial for its selective expression activity, especially for medical approaches as developed here against glioblastoma with highly vulnerable healthy brain tissue in direct contact to the tumor. Although knowledge about the exact functionality of IRES molecules is still incomplete, recent literature indicates that IRES secondary structures are rather flexible and can induce ribosomal dependent translation precisely because of this flexibility^37^. The IRES blockers (IB) we use represent complementary sequence motifs to IRES sequences and block the IRES functionality by attachment. Further investigations will show whether this function is achieved by altering IRES secondary structure or mainly by sterically blocking the IRES-ribosome interaction. However, our results on different IRES blockers suggest a combined effect since on the one hand all seRNA constructs with differing IB could reinduce IRES-dependent translation after activation in target cells. On the other hand, reduced activation levels of IB3 seRNA which interferes with the remarkably stable and functionally important IRES pseudoknot region^38, 39^, also suggests functional refolding of certain IRES domains after activation of seRNAs. Partially perturbed refolding might also explain why transfection efficiencies of seRNAs in target cells are generally slightly lower than those of seRNA_Δ3-5_ constructs.

To allow cell-type specific activation, RNA double strands between seRNA antisense and cell type-specific RNA must occur. Sense-antisense interactions are a common, natural regulatory mechanism that acts at different levels and can regulate both, the transcription itself and the amount of free sense-RNA molecules^24, 40^. Upon formation, RNA-RNA double-strands are preferentially recognized by RNase III family members^41^. However, although with a 25-fold preference for RNA/DNA hybrids^42^, RNase H1 can also bind and digest to some extent RNA/RNA double strand duplexes^43, 44^. This induces further processing after RNA cleavage or causes complete Xrn1-dependent degradation of RNA^45, 46^. For seRNA molecules we also could demonstrate a small but significant influence of RNase H on recognition and primary cleavage. Further analysis will show whether RNase III family members are the main proteins responsible for the activation of seRNAs. Unfortunately, proper inhibitors are missing and only expression of bacterial YmdB has been described to modulate RNase III activity^47^. It will also be interesting to see which length of antisense sequences used in the seRNA can be shortened and whether mismatches are tolerated. Analyses for RNase H and RNase III indicate that even much shorter antisense sequences should be sufficient to induce the activation of seRNAs^48, 49^. However, it is important to note that the function principle of seRNA is entirely different from those of siRNAs and miRNAs, since the latter interfere with cell specific mRNA to induce its degradation^50^. In contrast seRNAs use such sense-antisense interaction for intrinsic molecule activation for subsequent translation of the effector coding sequence.

RNase-induced complete degradation of seRNA molecules upon activation in target cells is inhibited by RNA sequence domains of the dengue virus. These sequences form specific secondary structures to inhibit further degradation by presumably altering the tertiary structure of the Xrn1 exonuclease to naturally induce the formation of a disease-related small flaviviral RNA^51, 52^. The identical function of this sequence is also conserved in seRNA molecules and thus allows degradation of the 5′-upstream sequences including the IRES blocker, but stabilizes the IRES and effector sequences for cell-type-specific activation of translation.

In non-target cells, seRNA remains inactive. Besides using IRES-blocker sequences CAP-dependent translational initiation also had to be blocked for full-length seRNAs. We have achieved this by incorporating upstream open reading frames (uORFs) in domains 2 to 6. The general mode of action of uORFs was already identified more than 35 years ago for yeast GCN4 mRNA^26^ and is also demonstrated for animal transcripts^53^. Depending on the surrounding sequence of a stop codon, the length of the uORF itself as well as the distance between the reading frames, they regulate not only the translation initiation at the start codon of the next ORF but also the partial loss of ribosomal subunits from the mRNA itself^22, 54^. For our seRNA constructs, up to 6 uORFs were used. At the same time, it was ensured that the last uORF overlapped in a different reading frame with the start codon of the effector, thus efficiently suppressing CAP-dependent translation in non-target cells. We assume that further optimization of uORFs can ultimately block the already only very low leakage in some non-target cells and thus ensure the most efficient translation suppression during the lifetime of full-length seRNAs. Innate immune responses can largely hamper the effectiveness of various medications. Especially in viral systems, pathogen-associated molecular patterns represent optimal recognition motives for intracellular and extracellular pattern recognition receptors (PRRs). Thereby, particular viral sequence motifs, secondary structures and especially dsRNAs are efficiently recognized mainly by toll-like and RIG1-like receptors and converted into an antiviral immune response^55, 56^. Our data indicate that intracellularly formed seRNAs are not recognized by PRRs, no matter if sense-antisense interaction is taking place in target cells or not. Although in vivo conditions are certainly a lot more complex and delivery-induced immune responses might be necessary, the lack of IL-6 expression indicates an extensive medical applicability. In addition, although seRNA molecules perform their function at the RNA level, they do not necessarily have to be delivered as RNA. This results in the broadest possible range of delivery options.

For proof of biotechnological and medical applicability we equipped our seRNAs with a continuously active form of caspase3^57^ as effector and tested targeted expression in U87 glioblastoma cells and in an in vivo mouse glioblastoma model. Glioblastoma defines one of the most aggressive grade 4 tumors with median survival times of just 15 months even in patients who receive best-practice treatment^58^. Different treatment approaches have been developed to enter preclinical and clinical phases including siRNA, mRNA, and small molecules^59, 60^. Basically all approaches failed for different reasons. However, the decisive factors were the lack of suitable glioblastoma surface markers and the resulting lack of glioblastoma-specific treatment in a highly sensitive tissue environment. At this time, we also cannot guarantee that seRNA will mark a breakthrough in glioblastoma treatment. However, our seRNA approach impressively shows that intracellular molecules can no longer be used only as diagnostic markers but, for the first time, function as therapeutic markers to facilitate the targeted formation of therapeutically highly active proteins exclusively in cancer cells. Such specific activation was detectable in cell culture and glioblastoma mice with massive tumor reductions within days after a single treatment with anti-glioblastoma seRNAs. Parallel rigorous scoring of glioblastoma mice during treatment did not detect any behavioral or neuronal deficits, supporting high biocompatibility and tumor-specific activation of seRNAs. Further experiments will show if seRNA drug candidates against glioblastoma will also be functional against older tumors or can be used in humans. However, due to the simple modular system of seRNAs, it does not require much imagination to develop further applications in the fields of oncology, virology or gene therapy.

## Methods

### seRNA construction

pmCherryC1 (Addgene) was used as a backbone plasmid for seRNA construction. The human keratin 13 antisense fragment was amplified from plasmid HK13deltaT.eGFP (provided by Reinhard Windoffer, Aachen) with primers P1 and P2 (see supplementary table 1). IRES blockers were implemented with primers P3 and P4 (IRES blocker 1, IB1), P5 and P4 (IB2), and P6 and P4 (IB3). RNase inhibitor region was generated according to the Eurofins Genomics DNA RNA oligonucleotide synthesis protocol with primers P7 and P8 (part 1) and P9 and P10 (part 2) (see supplementary table 1). EMCV-IRES was isolated from pFR_wt (Prof. Palmenberg WI, USA) with primers P11 and P12. The constitutively active version of caspase 3 was published before^57^ and cloned as described. All primer sequences including restriction sites used for cloning are indicated in supplementary table 1. Primers were ordered from Eurofins Genomics.

### RNA secondary and 3-dimensional structure prediction

Influences of IRES blockers IB1-IB3 on seRNA secondary structure were predicted using RNAfold WebServer application (ViennaRNA package, version 2.4.13). Secondary structure predictions were based on minimum free energy (MFE) and partition function. Isolated base pairs were avoided. For 3D RNA structure prediction, 3dRNA v2.0 was used in optimization and slow assembly mode. Predicted 3D structures were transferred in dot-bracket format to the Chimera 1.14 software for 3D animations, image generation and 3D structure superimposition.

### Cell culture

All cell types were cultivated at 37°C in a humidified atmosphere containing 5% CO_2_. Human foreskin fibroblasts (HFF, ATCC) were cultivated in DMEM Glutamax medium (Gibco) supplemented with 10% FBS (Gibco) + 1x PenStrep (ThermoFisher). To cultivate the breast cancer cell line MCF-7, RPMI 1640 GlutaMax medium (Gibco) was used. Medium was supplemented with 10% FBS, 1x PenStrep, 1x NEAA MEM (Sigma Aldrich), 1x sodium pyruvat (ThermoFisher) and 10 μg/ml insulin (Sigma Aldrich). U87MG glioblastoma cells (ATCC) were cultivated in MEM medium supplemented with 10% FBS, 1x PenStrep, 1x NEAA MEM, and 1x L-glutamine (ThermoFisher). For primary cortical neurons from rat embryo neurobasal medium supplemented with 2% B-27 supplement, 0.1% gentamicin reagent solution and 0.25% GlutaMax was used. For co-culture experiments of U87 and primary cortical neurons, and U87 sphere formation, a 1:1:1 mixture of RPMI GlutaMax medium (supplemented with 10% FBS and 1x L-glutamine), U87MG medium and primary cortical neuron medium was used. Cells were seeded 24 h before plasmid transfer on 24-well plates (BD Falcon; Thermo Scientific, USA) for microscopy, flow cytometry, and qRT-analysis. Primary cortical neurons were isolated as described previously^61^.

### Transfection (in vitro)

For seRNA encoding plasmid transfer, Lipofectamine 3000 was used with 1 μg plasmid DNA. Transfection was stopped after 4 h by replacing with fresh medium. Primary cortical neurons were treated by nucleofection using the Amaxa nucleofector kit for neurons and Amaxa program G-013. Human foreskin fibroblasts were treated with Amaxa nucleofector kit [Kit NHDF, Amaxa program U-020 NHDF human neonatal] with 2×10^6^ cells and 2 μg plasmid DNA.

### RNase inhibition

For RNAse H inhibition, cells were incubated with 10 μM of (Z)-5-(3,4-dihydroxybenzylidene)-3-(4-hydroxyphenyl)thiazolidine-2,4-dione (Life Chemicals, Canada) in the medium for 3 h. After washing, cells were transfected with appropriate seRNA plasmids and incubated for additional times as indicated before crude RNA isolation.

### Coculture of primary cortical neurons and U87 glioblastoma

To cultivate primary cortical neurons and U87 glioblastoma cells, silicone rubber inserts with 1 mm diameter circular cut-outs were produced. U87 cells were seeded in these holes, cultivated overnight, and transfected with Lipofectamine 3000 as described above. Subsequently, PDMS stamps were removed and primary cortical neurons were seeded on top. These cells were previously treated by nucleofection to transfer corresponding seRNA or control constructs.

### U87 sphere formation

To induce sphere formation, U87 cells were seeded with 200,000 cells / 24 well plate (BD Falcon) in triple medium as indicated above. Cells were transfected with seRNA plasmids either 24 h (effect on tumor formation) or 5 days after seeding when spheres were fully developed.

### Immunocytochemistry

For immunohistochemistry cells were cultivated on 35 mm glass bottom Petri dishes. After cultivation for 24 h, cells were washed twice with PBS (2 ml prewarmed) for 5 min at 37°C. Afterwards, cells were fixed with 250 μl methanol for 5 minutes at room temperature (RT). Cells were fixed with 250 μl pure acetone for 20 seconds and dried. For rehydration, 500 μl of PBS were added to each sample. All samples were blocked with 5% milk powder in PBS for 1 hour at RT. PanKeratine primary antibody (Progen 10550, rabbit, polyclonal) was added 1:500 in 1% milk powder-PBS and incubated overnight on a 2D shaker at 4°C. Subsequently, samples were washed three times with 1 ml PBS for 10 minutes at RT. The secondary antibody (Alexa Flour 488 Goat Anti Rabbit, A11034) was incubated for 1 hour at RT in a 1:500 diluted antibody:PBS solution. Subsequently, samples were washed three times with 1 ml PBS for 10 minutes and twice with 1 ml H_2_O.

### Flow cytometry

Flow Cytometry (CytoFLEX S Flow Cytometer, Beckmann Coulter) determined eGFP-mRNA transfer efficiency and fluorescence intensity. Briefly, 24 h after transfection, the cells were trypsinized (0.05% trypsin-EDTA solution, Sigma Aldrich, USA) and centrifuged at 300g. Without fixation, cells were analyzed directly after resuspension in 250 μl of the corresponding culture medium. At least 10,000 cells were analyzed for granularity and size by forward scattering for each cell type. eGFP fluorescence intensities were measured using appropriate filter settings and gatings.

### Live-dead staining

Cell viability was analyzed by flow cytometry according to the manufacturer protocols using the LIVE/DEAD™ Fixable Red Dead cell stain kit (L34971) and violet Dead cell stain kit (L34950, Thermo Fisher) with appropriate filter settings. Cells in the supernatant were included by centrifugation also to include already detached cells.

### Cell proliferation analysis

Cell proliferation in U87 spheroids was analyzed following the product protocol using the CyQuant NF Proliferation Assay (Thermo Scientific). Fluorescence intensities were quantified using a plate reader (TECAN Austria GmbH, infinite M1000 PRO).

### Microscopy

Live cell analyses were performed in phase contrast and fluorescence 24 h after plasmid transfer at 37°C and 5% CO_2_ using a confocal laser scanning microscope (cLSM 710, Carl Zeiss Jena, Germany). The microscope was equipped with a 488 nm argon laser and appropriate filter settings to visualize eGFP and AlexaFluor488. All images of transfected cells showed a representative overview at the center of each substrate and were recorded with an EC Plan-Neofluar 10×/0.3 Ph1 objective (Carl Zeiss Jena). Immunofluorescence images were recorded with a Plan-Apochromat 20×/0.8 Ph2 objective (Carl Zeiss Jena). For all experiments, microscope settings were kept identically.

### qRT-PCR experiments

For qRT-PCR analyses, total RNA was isolated with the RNeasy Plus Mini kit (QIAGEN) at different time points after transfection. cDNA synthesis was performed using QuantiTect Reverse Transcription Kit (QIAGEN GmbH, Germany). cDNAs of interest were quantified using 0.1 μg of total cDNA using a StepOne Real-Time PCR System (Thermo Scientific) and TaqMan assay-specific primers (supplementary table 1b) and master mix (Thermo Scientific). As endogenous control, glyceraldehyde 3-phosphate dehydrogenase (GAPDH) was used. StepOne Software (version 2.0.2) was used for evaluation.

### Statistical analysis and data evaluation

Data are given as mean (s.d.). Analysis of variance (ANOVA) was used for multiple comparisons. A p-value of ≤ 0.05 was considered as significant. Significance levels of 0.05, 0.01 and 0.001 were displayed with one to three stars.

### In vivo experiments

#### Animal experiments

The State Agency for Nature, Environment and Consumer Protection (Landesamt für Natur, Umwelt und Verbraucherschutz North Rhine-Westphalia; number 81-02.04.2019.A396) approved all animal experiments, which were conducted under the national Law for Animal Protection and European regulations and guidelines. Nude male immunodeficient mice (MRI-Foxn1nu/nu; Charles River Laboratories, France) were housed in individually ventilated cages in a specific pathogen free environment in (Tecniplast®, Blueline). Food and water were sterilized and provided ad libitum.

#### Tumor cell preparation

Human glioblastoma U87 cells were used for tumor cell implantation in mice when a confluency of 80% was reached. A cell suspension of 10^6^ U87 cells was created and stored in a 1 ml Eppendorf tube on ice until use. Storage time was limited to a maximum of six hours.

#### Surgery

30 min prior to surgery, mice received Carporfen 4 mg/ subcutaneously (Rimadyl®, Zoetis). Anaesthesia was induced by isoflurane inhalation. Using a microinjector (SMARTouch, UMP3, WPI) and a microsyringe (NanoFil 10μl, WPI), 2 × 10^5^ U87 cells were injected into the striatum of the right hemisphere, at the stereotaxic coordinates 2.4 mm lateral, 0.5 mm rostral, and 4 mm ventral of bregma. The syringe was left in place for five minutes and then removed slowly (1 mm/2 min). Additional Carprofen was given for two more days after surgery. Daily scoring of the animals followed to quantify severity levels.

Two days after tumor cell injection, mice received microinjection of either the therapeutic caspase-expressing seRNA-construct (n=10), the control construct expressing eGFP only (seRNA-eGFP, n=8) or sodium chloride (NaCl, n=2), using the same stereotaxic coordinates described above. The total number of indicated animals were treated in 4 independent experiments. RNA constructs were prepared freshly for each animal and used instantly.

#### Magnetic Resonance Imaging (MRI)

MRI was performed on treated mice on day three, seven and fourteen after transfer of seRNA or sodium chloride. MRI was performed on either a 9.4 Tesla scanner (BioSpec 94/20; Bruker BioSpi), using a 1H quadrature cryogenic cooled surface coil (CryoProbe, Bruker BioSpin) or a 3 Tesla MRI system (Achieva®, Philips Healthcare, Best) with an 8 Channel Volumetric Rat Array (Rapid Biomedical GmbH, Rimpar). Mice were anaesthetized with isoflurane and vital parameters were controlled by a custom-made setup (Medres; Cologne). After an initial localizer sequence, T2-weighted images were obtained. On the 9.4T Bruker scanner, we used following parameters: imaging method = RARE, RARE-factor = 8, repetition time = 5500 ms, echo time = 10.833 ms, acquisition matrix = 256 × 256; slice thickness = 0.5 mm, and number of averages = 2. For the 3 Tesla scanner we worked with following parameters: Slice thickness 0.5 mm, fast imaging mode = TSE, TSE factor = 5, acquisition matrix = 132 × 130, echo time = 37 ms, repetition time = 2794 ms. With both MRI system, 28 slices were scanned.

#### Histology

Mice were sacrificed by decapitation after the last MRI. Brains were harvested and stored at − 80°C until further analyses. Brains were cut coronally in 20 μm thick slices (Leica CM3050). To investigate proliferation, tissue was stained against Ki67 (1:500, Rabbit Polyclonal, ab15580, Abcam), using a fluorescent secondary antibody (1:500, Alexa-Fluor Goat anti-Rabbit 568nm IgG (H+L), Invitrogen, ThermoFisher). All nuclei were counter-stained with Hoechst (1:500, Hoechst 33342, Life Technologie). Images were analyzed with an inverted fluorescent microscope in phase contrast (Keyence BZ-9000E, Keyence), and representative pictures were acquired.

#### Image Analysis

Tumor volumes were quantified in vivo on MR-images using the software program VINCI 5.06 (Max Planck Institute for Neurological Research Cologne)^62^. To define tumor volumes, volumes of interest (VOIs) were defined in each animal and at every investigated time point.

#### Statistics

Calculations were performed using the software GraphPad Prism (version 9.1.2). Normal distribution was analyzed via D’Agostino & Pearson test. Group differences were assessed by one-way repeated measures analysis of variance (ANOVA) and performed in both groups for the factor time. Additionally, Greenhouse Geisser correction was conducted to control for violations of sphericity. Statistical significance was assumed at p < 5%. Results are shown in a box and whiskers plot, reporting the median, quartiles and extreme values as well as single values for every time point.

---

**Supplementary table 1a+b.**
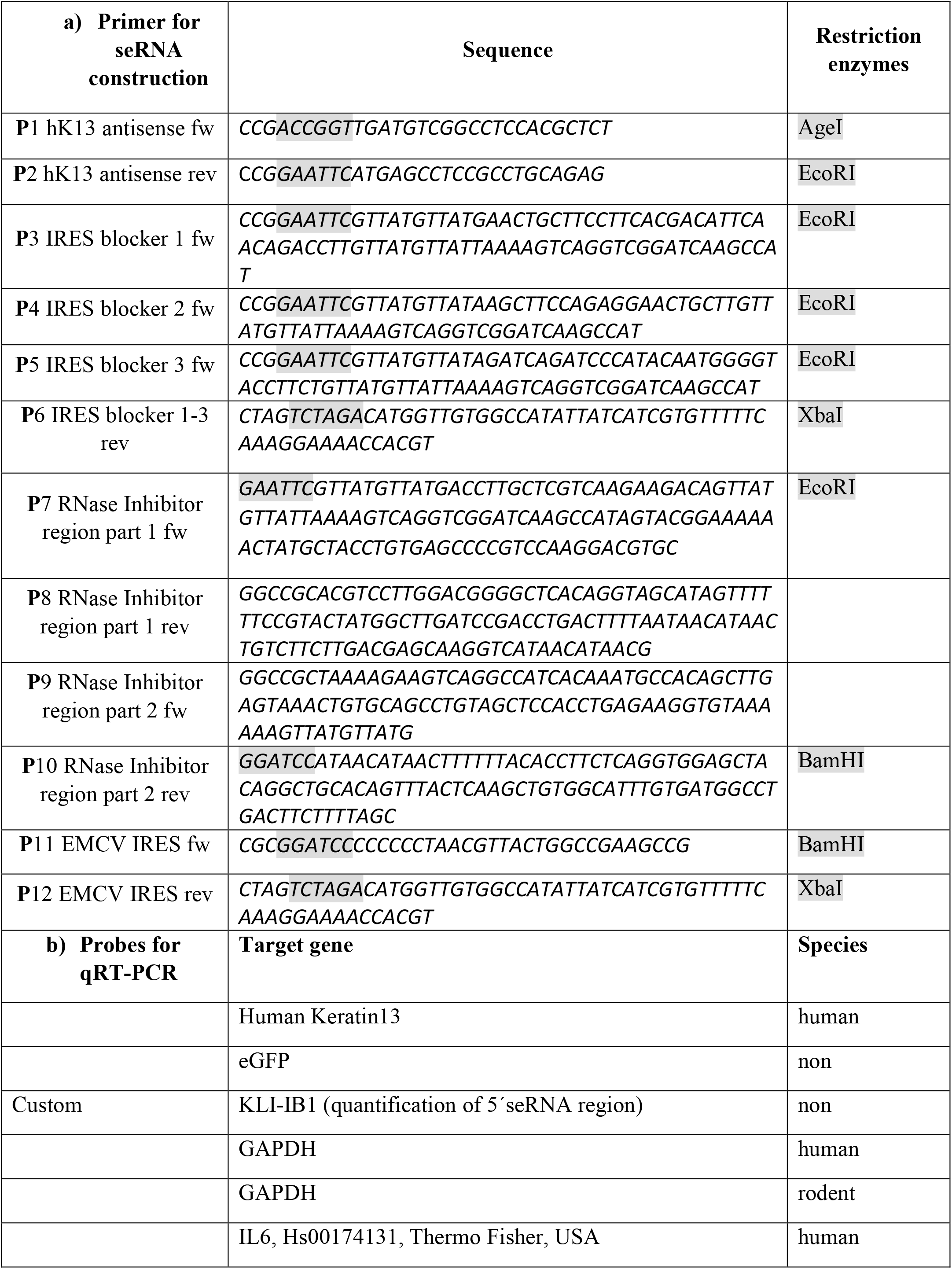
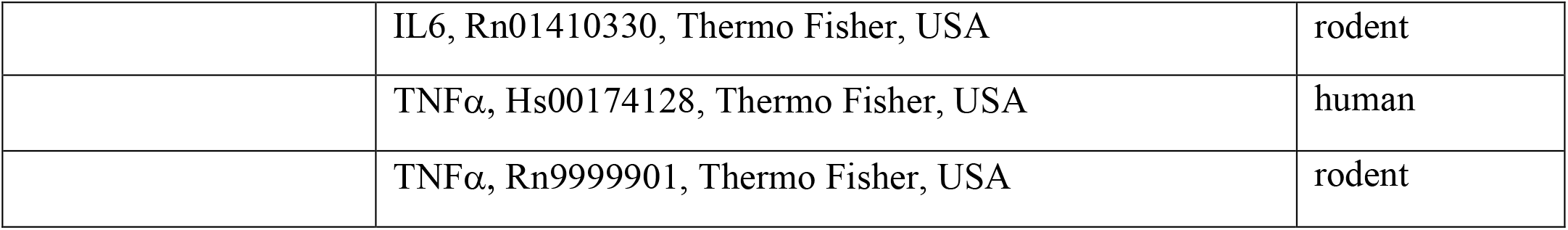

## Acknowledgment

We thank Saishreyas Iyer and Jennifer Gehlen for their experimental support. Special thanks goes to Prof. Windoffer (RWTH Aachen University, Germany) for providing the hKer13 plasmid, Prof. Palmenberg (Univ. of Wisconsin-Madison, WI, USA) for the EMCV-IRES plasmid and Prof. Langen (Research Centre Juelich, Germany) for providing the U87MG cells.

## Author contributions

BH developed the system, BH and MAR designed experiments; MH, SYS, HLW, CH, SG, NH and FR performed experiments and data analysis, DW supported MRI analyses, HB provided medical knowledge and verified therapeutic approach; BH, GRF, MH and RM wrote the paper. All authors read and approved the final manuscript.

